# Characterization and Variation of the Rhizosphere Fungal Community Structure of Cultivated Tetraploid Cotton

**DOI:** 10.1101/466912

**Authors:** Qinghua Qiao, Jingxia Zhang, Changle Ma, Furong Wang, Yu Chen, Chuanyun Zhang, Hui Zhang, Jun Zhang

**Affiliations:** Key Laboratory of Plant Stress Research, College of Life Sciences, Shandong Normal University, Jinan, 250014, China; Key Laboratory of Cotton Breeding and Cultivation in Huang-Huai-Hai Plain, Ministry of Agriculture, Cotton Research Center of Shandong Academy of Agricultural Sciences, Jinan, 250100, China

**Keywords:** cotton rhizosphere, fungal community, diversity, soil resource, developmental stage

## Abstract

Rhizosphere fungal communities exert important influential forces on plant growth and health. However, information on the dynamics of the rhizosphere fungal community structure of the worldwide economic crop, cotton (*Gossypium* spp.), is limited. Next-generation sequencing of nuclear ribosomal internal transcribed spacer-1 (ITS1) was used to characterize the rhizosphere fungal communities of worldwide cultivated tetraploid cotton using *G. hirsutum* cv. TM-1 (upland cotton) and *G. barbadense* cv. Hai 7124 (island cotton). Plants were grown in field soil (FS) that had been continuously cropped with cotton and nutrient-rich soil (NS) that had not been cropped. Fungal species richness, diversity, and community composition were analyzed and compared among soil resources, cotton genotypes, and developmental stages. We found that the fungal community structure between the rhizosphere and bulk soil of cotton were different and the rhizosphere fungal communities were significantly varied between FS and NS. These results suggest that cotton rhizosphere fungal community structure variation was primarily determined by the interaction of cotton roots with different soil resources. We also found that the community composition of cotton rhizosphere fungi varied significantly during different developmental stages, suggesting that developmental stages were also important factors in the dynamics of rhizosphere fungal communities for the varying dominant fungal genera of the rhizosphere. In addition, we also observed that fungal pathogens were clearly increased at certain developmental stages, suggesting a higher infection rate and a high incidence of corresponding soil-borne disease in each stage. This research illustrates the characteristics of cotton rhizosphere fungal communities and provides important information for understanding the potential influences of rhizosphere fungal communities on cotton growth and health.

## Introduction

Soil microorganisms are a critical component of agroecosystems and play key roles in agricultural ecosystems. The importance of mutual influence between microbial communities and agronomic practices is increasingly being recognized. The rhizosphere is the adjacent soil environment that the plant helps to create and where beneficial and pathogenic microorganisms exert major influential forces on plant growth and health [1]. Rhizosphere microorganisms were thought to be of great importance to plant health due to their involvement in such key processes as the formation of root architecture [2]; formation of soil characteristics [3]; decomposition of organic matter [4, 5]; decomposition and removal of toxins [6, 7]; defense against plant pathogenic microorganisms [2]; and cycling of carbon [8], nitrogen, phosphorus, and sulfur [9-12].

Soil fungi are a critical component of agroecosystems, and the rhizosphere fungal communities play important roles in plant growth and health. In turn, plants largely control rhizosphere fungi through the production of carbon- and energy-rich compounds and bioactive phytochemicals [13]. Some of the beneficial fungi are directly involved in the cycling of nutrients and function as an essential link to soil nutrient availability [14-17]. Some fungi are known for having biocontrol activity against pathogenic microorganisms [17, 18]. These fungi positively influence plant productivity by enhancing plant growth. However, certain rhizosphere fungal species or genera can also negatively influence plant productivity by causing disease, and pathogenic fungi are some of the most serious plant pathogens, for example, stalk rot disease of maize caused by *Fusarium* species [19], *Verticillium* wilt caused by *Verticillium nonalfalfae* on tree-of-heaven [20], and dry root rot caused by *Macrophomina phaseolina*, which affects many crops [21].

It is known that microbial diversity in soil is one of the major components determining soil health [22] and is believed to be one of the main drivers in disease suppression [22-25]. The composition of rhizosphere microbial communities is affected by soil, plant developmental stage, and many other factors [26-30]. Continuous cropping in agricultural production can cause crop yield reduction through soil quality degradation and aggravated plant diseases [31-33]. The fundamental reason for continuous cropping obstacles is related to disorders or deterioration of rhizosphere microorganisms (including rhizosphere fungi) [34, 35].

Cotton (*Gossypium* spp.) is the most important cash crop in the world and provides the most natural textile fibers of the world. Cotton production is threatened by soil-borne plant pathogens such as *Rhizoctonia* spp. [36], *Fusarium moniliforme* [37], *Alternaria alternata* [38], and *Verticillium dahliae* [39]. Understanding the dynamics of the rhizosphere fungal community structure of the worldwide cultivated tetraploid cotton with cotton cultivars in different developmental stages will not only provide basic information on the dynamics of cotton rhizosphere fungal community structure but also help lay a foundation for understanding the mutual influence between rhizosphere fungal communities and the plant health of cotton. Knox et *al*. showed that rhizosphere microbial diversity in cotton is significantly influenced by cultivar type in the field [40]. However, systematic studies on the rhizosphere fungal community structure of cultivated tetraploid cotton are still lacking.

This study characterized the rhizosphere fungal community dynamics across cotton developmental stage growth using two cotton cultivars in continuously mono-cropped cotton field soils (FS) and nutrient-rich soil (NS) that had not been cropped. Our work lays the foundation for more research on cotton rhizosphere fungal communities and may provide insight into further dissection of the structure of rhizosphere fungal communities, which might exert major influential forces on plant growth and health in the agricultural production of cotton.

## Materials and methods

### Plants and soil

Two cultivars of cultivated allotetraploid *Gossypium* species, *G. hirsutum* cv. TM-1 (upland cotton) and *G. barbadense* cv. Hai 7124 (island cotton with higher disease resistance than upland cotton) were planted in two types of soils FS and NS.

The FS was obtained from 15 to 30 cm below the soil surface in a field that has been continuously planted with cotton for several decades at the Experiment Station of Cotton Research Center of Shandong Academy of Agricultural Sciences (Linqing County, Shandong Province, 36°81′N, and 115.71°13′E), and the NS, which was not influenced by cotton and any other plants, was purchased from Feng Yuan Science and Technology Ltd. (Jinan, China). All visible biota (e.g., weeds, twigs, worms, and insects) were removed, and the soil was then crushed and sifted through a sterile 2 mm sieve. Because the sieved soil drained poorly and was difficult to sample, we mixed sterile sand into the treatment soils at a soil:sand ratio of 2:1 following Lundberg *et al*. [41].

All plants were grown under the same environmental conditions. Samples were collected at the seedling, budding, and flowering stages. Detailed information about the material and methods were described in our previous report [42].

### Greenhouse plant management

Cotton seeds were delinted by sulfuric acid and then surface sterilized with 75% ethanol for 15 min, followed by 30% H_2_O_2_ for 30 min and five rinses with sterile distilled water. The seeds were germinated by incubating at 28 °C in the dark for 2–3 days in petri dishes in which sterile paper was overlaid on 1% water agar. After germination, seedlings were transplanted into the treated soil and raised in a tissue culture room at 28 °C. Plants were moved to a bioclean greenhouse as soon as seedlings developed a second true leaf. The pots were watered every 3 days with sterile water. Control pots contained soil without a cotton plant.

### Sampling of cotton rhizosphere and bulk soil

Soil samples were harvested from 4 to 6 July 2015. Well-grown plant individuals in each developmental stage were selected for rhizosphere soil collection. We inverted each pot to remove the soil and plant and then gently shook the plant to remove the soil that did not adhere to the root surface. Rhizosphere soil consisted of ~1 mm of soil that tightly adhered to the root surface and was not easily shaken from the root. To separate the rhizosphere soil, roots were placed in a sterile flask with 50 ml of sterile phosphate buffered saline solution and stirred vigorously with sterile forceps. Samples at the interface or from an unnatural environment were avoided. After cleaning, the roots were removed, and the remaining soil solution was centrifuged for 15 min at 10,000 rpm. The supernatant was discarded to leave the soil fraction. These soil fractions were frozen using liquid nitrogen and stored at −80 °C. We also collected samples from unplanted pots from ~10 cm below the soil surface as bulk soil. There were three biological replicates for each soil treatment (rhizosphere samples of the two cultivars and bulk soil samples in FS and NS were collected at three developmental stages) for a total of 54 replicates.

### DNA extraction and detection

The DNA from each soil sample was extracted using the Omega D5625-02 Soil DNA Kit (Omega Biotek Inc., Norcross, GA, USA). DNA concentration and integrity were detected by a microplate reader (Qubit 3.0 Fluorometer; Thermo Fisher Scientific, Waltham, MA, USA) and agarose gel electrophoresis (PowerPac Basic164-5050 and Sub-Cell 96, Bio-Rad Laboratories, Hercules, CA, USA). DNA information for each sample is listed in the Supplementary materials

### Preparation of libraries and sequencing

All suitable DNA samples were submitted to BGI Tech Solutions Co., Ltd. (Shenzhen, China) to construct a sequencing library. DNA from 54 soil samples was amplified and sequenced using the Illumina MiSeq platform (Illumina, San Diego, CA, USA). Further details on the subsequent bioinformatics analysis of the sequencing data are listed in the Supplementary materials and methods.

### Data analysis

OTU Venn diagram: The presence or absence of operational taxonomic units (OTUs) was determined for each soil sample, and the common and specific OTU IDs were summarized. A Venn diagram was constructed using the package VennDiagram in R (v 3.0.3).

Species Annotation: The tag number of each phylum in different soil samples was summarized in a histogram, and all data were used to construct a histogram using R.

α-diversity analysis: The species accumulation curves of observed species (Sobs), Chao, Abundance Based Coverage Estimator (ACE), Shannon, and Simpson indices were calculated using the software Mothur (v 1.31.2). The calculation formula of each index can be found at http://www.mothur.org/wiki/Calculators.

β-diversity analysis: β-diversity was analyzed using the software QIIME (v 1.80). Normalization was performed to control for sequencing depth differences in different samples. Sequences were extracted randomly according to the minimum sequence number of all samples to generate a new ‘OTU table biom’ file. Then, the β-diversity distance was calculated based on the ‘OTU table biom’ file. The β-diversity heat map was drawn by the ‘aheatmap’ function in the ‘NMF’ package of R.

Contribution of each factor: The Bray–Curtis dissimilarity analysis and the information entropy method were used to measure the contribution of the different factors to variability between samples. We then conducted an analysis of variance by the function aov in the R package. Interaction between each of the two factors was considered. For each factor, the contribution rate to fungal community variance was calculated as the mean square of the factor divided by the sum of the mean square of all factors.

## Results

Fungal communities were characterized by next-generation sequencing of nuclear ribosomal internal transcribed spacer-1. A total of 5,032,042 high-quality reads were obtained with a median read count of 93,186 per sample (range: 51,752–244,354) (Supplementary Table S1). The high-quality reads were clustered into 1,298 microbial OTUs at 97% similarity after the removal of OTUs that were unassigned or not assigned to the target species.

### Fungal communities in bulk soils of FS and NS

Ascomycota, Basidiomycota, and Zygomycota were the most common fungal phyla in both continuously cropped field soil (FS) and nutrient-rich soil (NS) treatments, accounting for 59.01–95.81% of all fungal communities (Supplementary Table S2; Supplementary Fig. 1). Excluding unclassified orders (19.39–60.96% of total fungal communities), in both soils, Eurotiales and Hypocreales were dominant in Ascomycota, and Mortierellales was dominant in Zygomycota. The dominant orders of Basidiomycota in FS were Cystofilobasidiales and Sporidiobolales, whereas Thelephorales and Agaricales were dominant in NS.

The differences in fungal communities between the FS and NS soils at the genus level were significant. The relative abundance of some fungal genera, such as *Penicillium*, *Gliomastix*, and *Engyodontium*, was significantly lower in FS than in NS (*P* < 0.05), whereas the relative abundance of some fungal genera, such as *Pseudozyma*, *Panaeolus*, and *Lecanicillium* in FS was slightly, but not significantly, higher than in NS (Supplementary Table S2).

### Fungal communities of cotton rhizosphere in FS and NS

Ascomycota, Basidiomycota, and Zygomycota were the dominant phyla in the rhizosphere fungal communities, accounting for approximately 33.45–88.51% of the total fungal communities in NS (11.48–66.15% were unclassified) and 85.18–93.88% of the total fungal communities in FS (6.03–14.65% were unclassified) (Fig. 1; Supplementary Table S3; Supplementary Figs. 2, 3). Ascomycota was negatively selected in the rhizosphere in NS but was enriched in the rhizosphere in FS (Fig. 1; Supplementary Table S2–4). The dominant orders of Ascomycota and Zygomycota in the rhizosphere were the same as those in bulk soil (Supplementary Table S3). However, the dominant orders of Basidiomycota in bulk soil from the FS rhizosphere samples were Agaricales and Auriculariales, whereas Sporidiobolales and Agaricales dominated in bulk soil from the NS rhizosphere samples (Supplementary Table S3).

**Fig 1.**
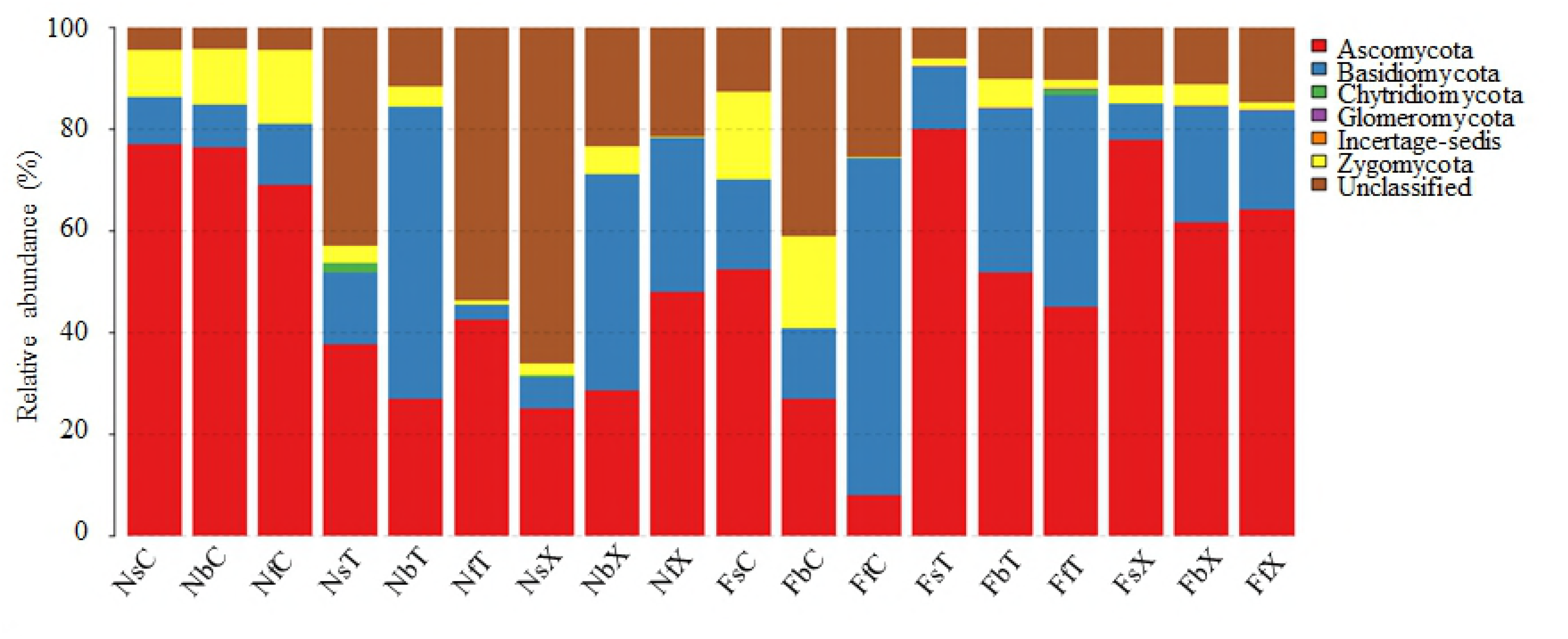
Relative abundance of the fungal community in all treatments. Two types of soils: nutrient-rich soil (N) and continuous cropping field soil (F). Three cotton plant developmental stages: seedling stage (s), budding stage (b), and flowering stage (f). Two cultivated species: upland cotton (*G. hirsutum* L. cv TM-1) (T) and sea island cotton (*G. barbadense* L. cv Hai7124) (X) and control pots (C) that lacked cotton plants. Each sample was labeled by a three-letter code, such as NsT, which indicates seedlings of sea island cotton grown in nutrient-rich soil.

The number of OTUs in the FS rhizosphere (205.33 ± 22.47) was higher than in FS bulk soil (140.67 ± 28.61), whereas in the NS rhizosphere (146.44 ± 40.22), the OTUs were lower than in NS bulk soil (181.11 ± 20.37) (Supplementary Table S5). The α-diversity of fungi was significantly higher in the FS rhizosphere than in FS bulk soil (*P* < 0.05); however, it was significantly lower in the NS rhizosphere than in the corresponding bulk soil (*P* < 0.05). Bulk soil α-diversity of fungi was higher in NS than in FS (*P* < 0.05), but rhizosphere fungal α-diversity was lower in NS than in FS (*P* < 0.05; Fig. 2; Supplementary Table S5).

**Fig 2.**
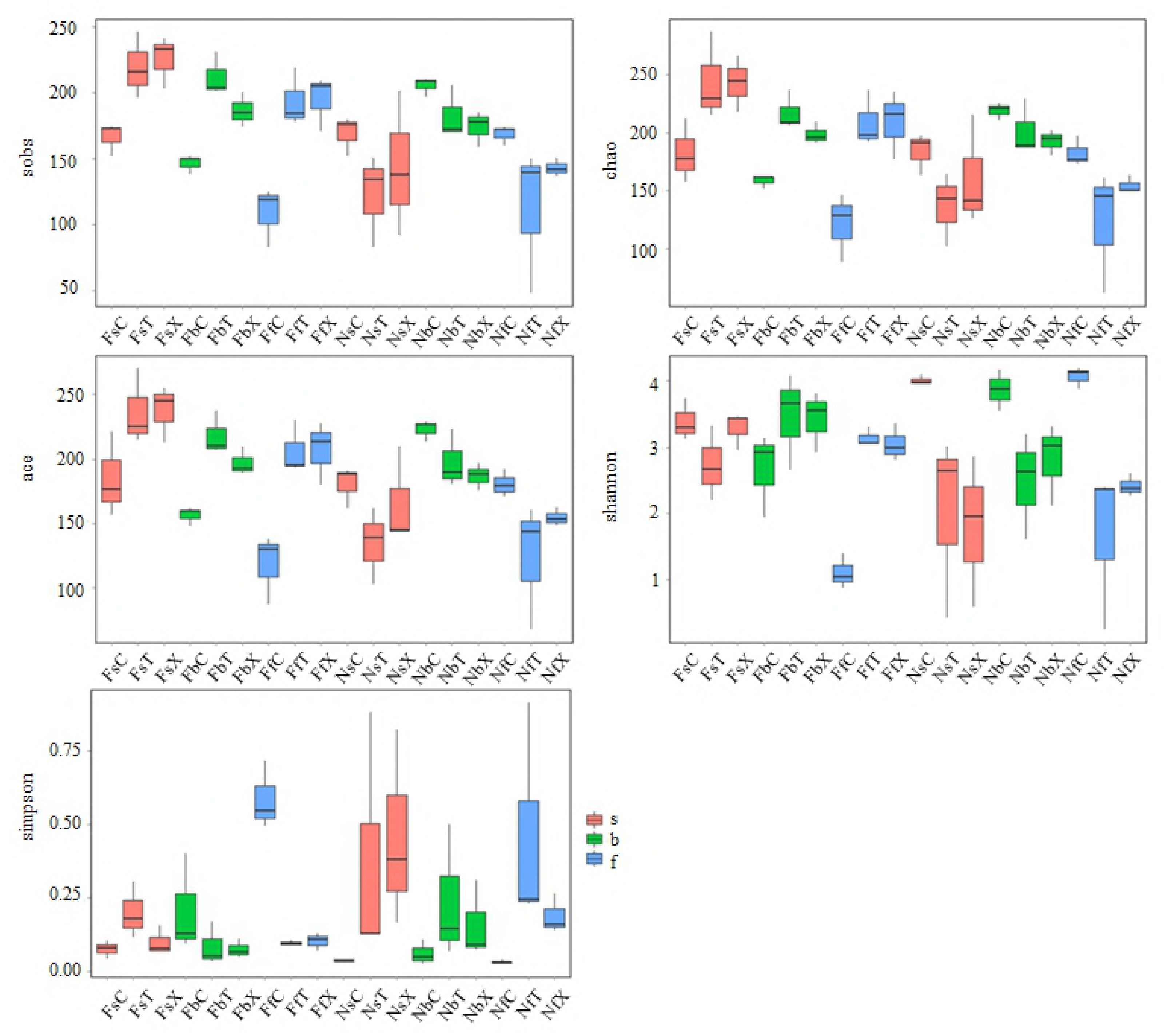
The α-diversity of rhizosphere fungi. From left to right and from top to bottom, box plots are Sob, Chao, ACE, Shannon, and Simpson indices.

Fungal genera compositions that were enriched or negatively selected in the rhizosphere differed between different soils (Supplementary Table S6; Supplementary Table S7). For example, in NS, the relative abundance of *Mortierella*, *Gliomastix*, and *Engyodontium* was significantly higher in bulk soil compared with rhizosphere soil, where it was much lower or almost undetectable (*P* < 0.05; Supplementary Table S8). In contrast, the relative abundance of *Rhodosporidium* and *Trichoderma* in NS rhizosphere soil was higher than in the respective bulk soil, where it was lower or almost undetectable (*P* < 0.05; Supplementary Table S8). In FS, the relative abundance of *Mortierella*, *Guehomyces*, and *Fusarium* was higher in bulk soil than in rhizosphere soil, where it was lower or undetectable (*P* > 0.05; Supplementary Table S9). The relative abundance of Penicillium, Alternaria, and Preussia was higher in FS rhizosphere soil than in bulk soil, where these genera were almost undetectable (*P* < 0.05; Supplementary Table S9). The abundance of other rhizosphere fungal genera was highly variable and differed between soils. Comparisons of fungal genera whose relative abundance changed inversely in different soils between the rhizosphere and corresponding bulk soil are listed in Table 1.

**Table 1.** Fungal genera that were affected inversely by cotton root in two soil resources.

| Genus | Relative abundance<br>in field soil (mean) |  |  | Relative abundance in<br>nutrient soil (mean) |  |  |
| --- | --- | --- | --- | --- | --- | --- |
|  | Control | Rhizosphere |  | Control | Rhizosphere |  |
| *Paraconiothyrium* | 0.21 | 0.00 | - | 0.00 | 0.10 | + |
| *Rhodosporidium* | 1.598 | 0.01 | - | 0.00 | 3.45 | + |
| *Mrakia* | 0.50 | 0.00 | - | 0.00 | 0.12 | + |
| *Arnium* | 0.28 | 0.00 | - | 0.00 | 0.08 | + |
| *Pseudeurotium* | 0.77 | 0.00 | - | 0.00 | 0.02 | + |
| *Kurtzmanomyces* | 0.17 | 0.00 | - | 0.00 | 0.02 | + |
| *Tomentella* | 0.05 | 0.10 | + | 0.45 | 0.22 | - |
| *Wardomyces* | 0.00 | 0.13 | + | 0.58 | 0.00 | - |
| *Chrysosporium* | 0.01 | 0.16 | + | 0.94 | 0.00 | - |
| *Retroconis* | 0.00 | 0.21 | + | 1.71 | 0.00 | - |
| *Nectria* | 0.00 | 0.22 | + | 1.07 | 0.00 | - |
| *Engyodontium* | 0.08 | 0.50 | + | 3.31 | 0.07 | - |
| *Gliomastix* | 0.00 | 0.41 | + | 2.85 | 0.00 | - |
| *Alternaria* | 0.17 | 0.66 | + | 0.88 | 0.46 | - |
| *Preussia* | 0.00 | 0.89 | + | 2.09 | 0.03 | - |
| *Penicillium* | 0.71 | 14.06 | + | 10.51 | 4.11 | - |

“+” denotes fungi with higher relative abundance in rhizosphere soil than in bulk soil, “-” denotes fungi with lower relative abundance in rhizosphere soil than in bulk soil; $P < 0.05$

### Variation in rhizosphere fungal communities at different plant developmental stages

In FS, the number of stage-specific OTUs was highest in the seedling stage and decreased gradually through development: upland cotton (T): 90 (seedling stage), 76 (budding stage), and 83 (flowering stage); island cotton (X): 121 (seedling stage), 53 (budding stage), and 48 (flowering stage). In NS, the number of stage-specific OTUs was highest in the budding stage (T: 71, 139, 85; X: 112, 138, 82). In addition, the number of overlapping OTUs in the seedling and budding stages was higher than that in the budding and flowering stages in both FS and NS soil treatments. The number of overlapping OTUs in all three developmental stages was higher in FS than in NS (Supplementary Fig. 4).

Analysis of α-diversity indicated that in FS, the Sobs, Chao, and ACE indices were higher in the cotton rhizosphere fungal communities during all three developmental stages compared with bulk soil. The Sobs index decreased gradually from the seedling to the flowering stage in bulk soil, but no significant difference was found in the rhizosphere sample between different developmental stages (except for the difference between the seedling stage and budding stage in the rhizosphere of island cotton) (*P* < 0.05; Fig. 2; Supplementary Table S5). In NS, the rhizosphere harbored a fungal community of higher α-diversity than bulk soil. We compared the α-diversity of different samples from NS to that of FS. The Sobs, Chao, and ACE indices indicated that the α-diversity of bulk soils from FS was generally lower than those from NS, but not significantly. In contrast, rhizosphere soils from FS were significantly higher than those from NS (*P < 0.05; Fig. 2; Supplementary Table S5)*.

Each developmental stage had dominant fungal genera found with high relative abundance. We determined the genera that had high relative abundance (relative abundance >0.5) in the different developmental stages. In the rhizosphere soils, Penicillium, *Fusarium*, *and Mortierella* in FS and *Penicillium*, *Fusarium*, and *Talaromyces* in NS presented a higher relative abundance in all three developmental stages. In addition, each developmental stage harbored the specific dominant rhizosphere fungal genera (Supplementary Table S10). The number of dominant genera was highest in the budding stage.

We also analyzed how the fungal community was affected by the presence of cotton. A large change was defined as a difference in relative abundance between rhizosphere and bulk soil that was >1 or <-1. The difference between rhizosphere and bulk soil fungal genera relative abundance differed at different developmental stages. We defined a genus for which relative abundance was greater in rhizosphere soil compared with bulk soil as an enriched fungal genus (EFG) and a genus for which abundance was lower in rhizosphere soil compared with bulk soil as a depleted fungal genus (DFG). EFGs were most abundant in the budding stage, whereas DFGs were most abundant in the seedling stage. The number of DFGs in NS was higher than in FS, in accordance with our finding that the α-diversity of fungal communities was higher in NS than in FS, and many fungi were depleted under the influence of cotton root (Supplementary Table S10).

We analyzed the β-diversity of the samples based on Bray–Curtis dissimilarity analysis. Cluster analysis indicated that samples from the same soil resources were clustered into one group (Fig. 3A). The β-diversity of different soils (mean Bray–Curtis: 0.97) was significantly higher than the β-diversity of different developmental stages (mean Bray–Curtis N: 0.66, F: 0.60) (*P* < 0.01; Supplementary Table S11; Fig. 3B). Statistical analyses were conducted to assess the contribution of each factor to the structure of the fungal community in the cotton rhizosphere and found that species-level soil factors contributed approximately 42.27% to the fungal community structure in the cotton rhizosphere, which was higher than other factors (*P* < 0.05; Supplementary Table S11).

**Fig 3.**
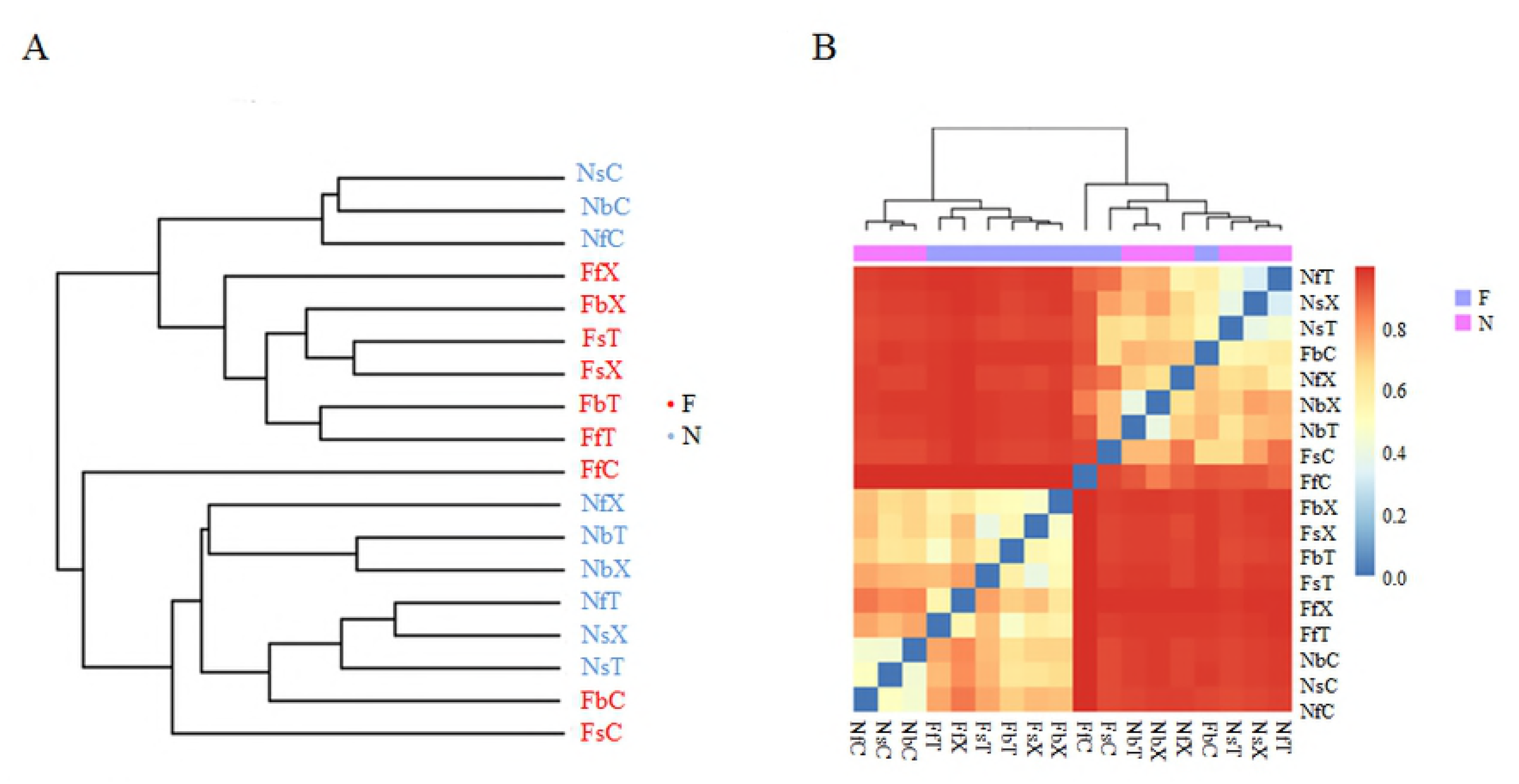
β-diversity analysis of different treatments. A: Cluster analysis of different treatments. B: Bray–Curtis distance analysis of different treatments.

### Potential pathogenic and phosphate-solubilizing fungi in the cotton rhizosphere

Pathogenic fungi were mainly distributed in the genera *Alternaria, Fusarium, Gibberella* [43], *Rhizoctonia, Thanatephorus* [44], and *Verticillium*. We analyzed the dynamics of those genera in different soils and found that the relative abundance of each genus in the rhizosphere was higher in bulk soil in pots containing FS but lower in pots containing NS (Supplementary Table S12). In addition, the relative abundance of these genera differed during different plant developmental stages. In FS, the greatest difference in the relative abundance between bulk soil and rhizosphere was present in *Alternaria* and *Rhizoctonia* at the seedling stage, and *Fusarium, Thanatephorus, Verticillium*, and *Gibberella* at the budding stage (Fig. 4; Supplementary Table S12). The rhizosphere relative abundance of *Fusarium* was lower than bulk soil at the seedling stage, and *Rhizoctonia* was lower than bulk soil at the budding stage. We conclude that in continuously cotton-cropped soil, those genera were suppressed by the cotton root at different stages. In NS, the relative abundance of most of these genera was lower in rhizosphere soil than in bulk soil, with the exception of the seedling stage for *Alternaria* and *Fusarium*, the budding stage for *Fusarium* and *Rhizoctonia* and the flowering stage for *Gibberella* (Fig. 4; Supplementary Table S12). Cotton growth in soil that had not been cropped might have a high infection rate at each stage by those genera. The difference in these genera between the two genotypes was not significant (Supplementary Table S12). The relative abundance of disease-associated fungal genera, with the exception of *Fusarium* (FS: 2.02–43.19; NS: 3.17–7.40), was higher in NS than in FS (*P* < 0.05), such as Verticillium (FS: 0.14–1.19; NS: 3.89–5.29) and Alternaria (FS: 0.04–0.85; NS: 2.01–3.27; Supplementary Table S2).

**Fig 4.**
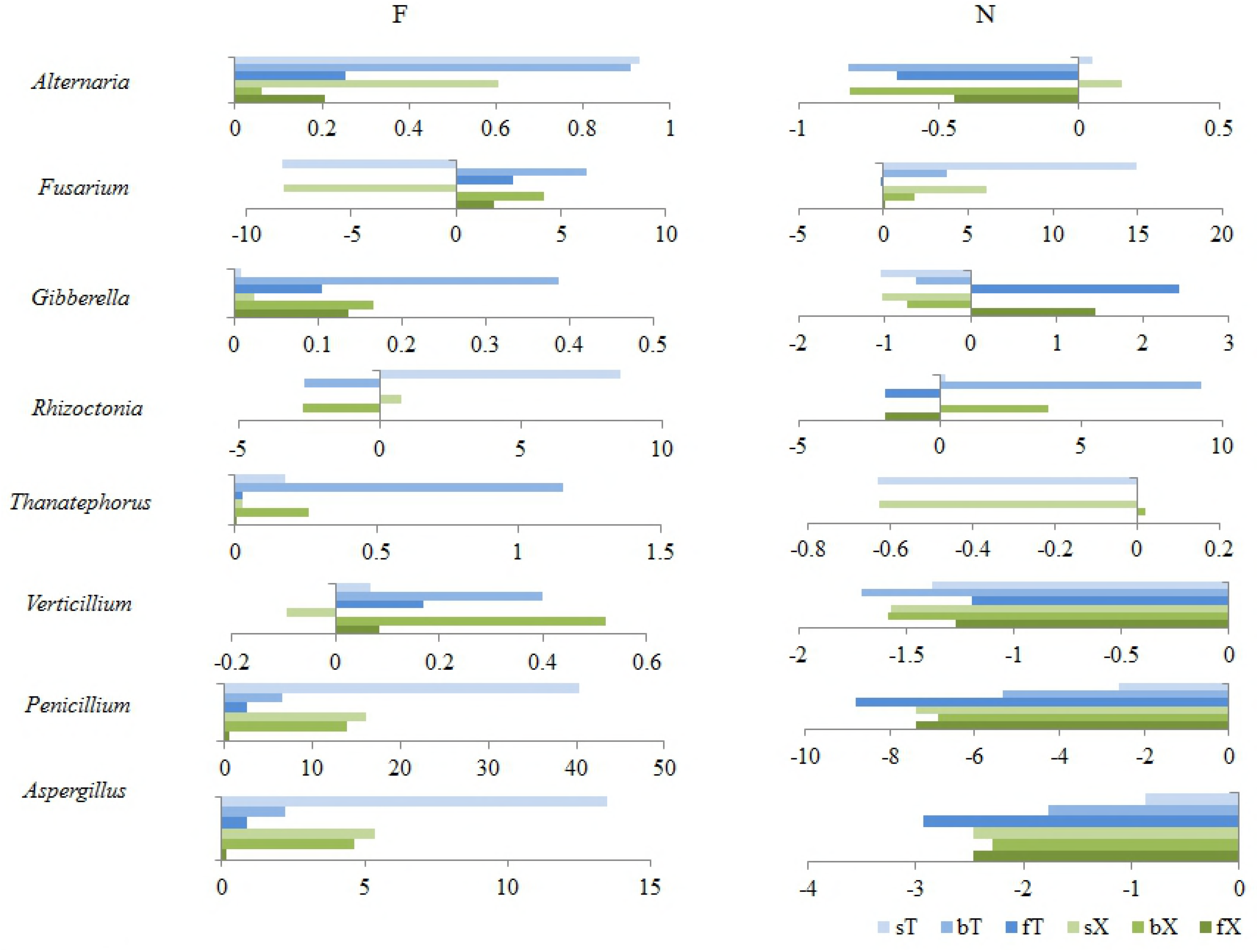
Variations of potential pathogenic and phosphate-solubilizing fungal genera. The X-axis shows different values of relative abundance between rhizosphere soils and bulk soils.

*Aspergillus* and *Penicillium*, the potential phosphate-solubilizing fungal genera, were detected in our research. In NS, the relative abundance of both fungal genera was lower in the rhizosphere than in bulk soil (*P* < 0.05). In FS, the relative abundance of the two fungal genera was higher in the rhizosphere than in bulk soil, but this difference was not statistically significant (*P > 0.05). In addition, the relative abundance of the two genera in rhizosphere soil was higher in* FS than in NS (*Aspergillus*: *P* < 0.01; *Penicillium*: *P* > 0.05; Supplementary Table S12).

## Discussion

### The difference in fungal community structure between the rhizosphere and bulk soil of cotton

Plant roots have a remarkable effect on the physical and chemical characteristics of soil, such as its structure and water retention [45-47]. The physical and chemical characteristics of the root-associated soil are important because they determine both the physiological aspects of root function, such as water and nutrient uptake, and the microbial activity that is most relevant to root growth [48-50]. Plant roots also release root exudates, volatile substances, border cells, and polymers into the soil environment and regulate the community structure of the rhizosphere microbiome through complex interactions with soil microorganisms [51-57], promoting the colonization of beneficial microorganisms and inhibiting the colonization of harmful microorganisms [58]. Many studies have confirmed the existence of differences in the microbial communities of rhizosphere soil and the surrounding bulk soil of *Arabidopsis*, rice and *Populus* [30, 41, 59].

In the present study, the dominant fungal phyla in the rhizosphere of the two cultivars of cultivated allotetraploid *Gossypium* species were Ascomycota, Basidiomycota, and Zygomycota, which is the same as that in bulk soils. The relative abundance of each phylum in rhizosphere soil differed from that of bulk soil to different degrees. Fungal communities influenced by cotton roots were mainly distributed in Basidiomycota. The dominant orders of Ascomycota and Zygomycota were the same in rhizosphere and bulk soils, but Basidiomycota was different. The dominant orders of Basidiomycota were Agaricales and Auriculariales in FS and Agaricales and Trechisporales in NS, which differed from that of bulk soil. Thus, we speculate that the soil-derived fungal community composition determines the rhizosphere fungal community of cotton, whereas cotton root affects the soil fungal community composition to a large extent. The β-diversity analysis and contribution analysis of each factor based on Bray–Curtis dissimilarity confirm the conclusion that the soil resource in this study is the main factor that determines the rhizosphere fungal community.

### Rhizosphere fungal communities varied in FS and NS

The characteristic of the soil itself is an important factor affecting the community structure of plant rhizosphere microorganisms. Moreover, the microorganism composition of soil is the main cause of variation in the community structure of the rhizosphere microbiome [60, 61]. In this study, significant differences were presented in rhizosphere fungal communities between different sources of soil. The difference was presented in two aspects: 1) The influences of cotton root on different fungal species were different. For example, in NS, the relative abundance of *Engyodontium*, *Mortierella*, and *Penicillium* was lower in pots containing cotton plants, whereas the relative abundance of *Clitopilus, Fusarium*, and *Rhodosporidium* was higher in pots containing cotton plants; 2) The influence of cotton root on some fungal communities differed substantially between NS and FS soil. For example, the relative abundance of *Mrakia*, *Rhodosporidium*, and *Talaromyces* in rhizosphere soil compared to bulk soil was higher in NS but lower in FS. This difference might be attributed to the different characteristics of the two soil resources. Thus, we conclude that the cotton rhizosphere fungal community structure variation was mainly determined by the interaction of cotton root with different sources of soil.

Rhizosphere microbial diversity can improve a plant’s resistance to soil-borne disease [17]. Previous studies have shown that continuous cropping can decrease the structural and functional diversity of the soil microbiome [62, 63]. In the present study, pots that did not contain plants had lower fungal α-diversity in FS than in NS, corroborating that long-term continuous cropping of cotton decreases fungal α-diversity, which in turn may be one of the important factors inducing continuous cotton-cropping obstacles. However, after planting with cotton, the fungal α-diversity of rhizosphere soils from FS was increased compared with bulk soil and higher than that of NS. We speculate that fungal communities in continuously cotton-cropped field soils might contain an abundance of fungi that are closely linked to cotton growth, nutrient absorption, and stress tolerance, and the functional limitation of such fungal communities is the main reason for continuous cotton-cropping obstacles.

### Developmental stages contributed to the variation of the fungal community in the cotton rhizosphere

Baudoin *et al*. proposed that the quantity and quality of root exudate input into the rhizosphere differ at different plant developmental stages, leading to differences in the composition of rhizosphere microbial communities between plant developmental stages [64]. Other studies have also demonstrated that rhizosphere microbes are significantly affected by the developmental stages of plants [65-69]. Our results indicate that the community composition of cotton rhizosphere fungi varied significantly during different developmental stages. The species richness of rhizosphere fungal communities was highest in the seedling stage in FS and in the budding stage in NS. In addition to the common dominant fungal genera of all three developmental stages, the rhizosphere fungal communities had a stage-specific dominant genus. The number of dominant genera and EFGs were the highest in the budding stage, which may be related to the plant requiring specific materials or releasing certain hormones into the soil during this stage.

### Alterations of potential pathogenic and phosphate-solubilizing fungal genera in the rhizosphere of cotton

Incidence rates of soil-borne disease are affected by many factors, such as the soil environment [70, 71], soil fungal community structure and function [17, 72, 73], relative abundance of pathogenic fungi, resistance of cotton cultivars, and developmental stage of cotton. Our results show that the relative abundance of disease-associated fungal genera in the bulk soil of FS and NS differed significantly. The relative abundance of potential pathogenic fungal genera (besides *Fusarium*) was lower in bulk soil of FS compared with that of NS. However, the relative abundance of these potentially pathogenic fungal genera in the rhizosphere was higher in FS and lower in NS compared with the corresponding bulk soil treatments.

The effect of cotton root on potentially pathogenic soil fungal genera also differed in different plant developmental stages. In FS, the relative abundance of *Alternaria* and *Rhizoctonia* at the seedling stage and *Fusarium, Gibberella*, *Thanatephorus*, and *Verticillium* at the budding stage in the cotton rhizosphere had the highest enrichment compared with bulk soil. In NS, the potentially pathogenic fungal genera were suppressed in rhizosphere soil, with the exception of the seedling stage for *Alternaria* and *Fusarium*, the budding stage for *Fusarium* and *Rhizoctonia* and the flowering stage for *Gibberella*. We speculate that potentially pathogenic fungal genera enriched in a developmental stage have a high infection rate of cotton root and thus cause a high incidence of soil-borne disease. The incidence rate was higher in FS than in NS and highest in the budding stage.

Diseases associated with fungal genera also differed by cotton genotype. Upland cotton (TM-1) was more susceptible to disease than island cotton (Hai7124), but this difference was not significant.

Fungi play an important role in the absorption and transformation of nutrients, especially phosphate-solubilizing fungi [14-16, 74]. Fungal species of *Aspergillus* and *Penicillium*, such as *Aspergillus tubingensis*, *Aspergillus niger* [75], *Aspergillus awamori, Penicillium citrinum* [15], *Penicillium albidum* [76], and *Penicillium oxalicum* [77], play an important role in phosphate solubility. We analyzed the dynamics of the two potential phosphate-solubilizing fungal genera. In cotton rhizosphere soils, the relative abundance of the two genera was higher in FS than in NS. This may be attributed to differences in physical and chemical properties and utilization of nutrient substances.

Our study provides insights into the structural variation of rhizosphere fungal communities under the influence of soil resources, developmental stage, and genotype, which might play key roles in cotton growth and health. The soil resources, cotton developmental stage, and cotton genotype all impacted cotton rhizosphere fungal community composition. The composition of the cotton rhizosphere fungal community was primarily determined by soil resources and regulated to a certain degree by plant developmental stage. A limited effect was found for the cotton genotype.

## Supporting information

**Supplementary Fig S1. Relative abundance of fungal phyla in bulk soil of both soils.**

**Supplementary Fig S2. Relative abundance of fungal phyla in the rhizosphere of cotton planted in field soil that has been continuously cotton-cropped.**

**Supplementary Fig S3. Relative abundance of fungal phyla in the rhizosphere of cotton planted in nutrient-rich soil.**

**Supplementary Fig S4. Total number of OTUs of specific and common fungi in different treatments.**

**Supplementary materials and methods S1**

**Supplementary Table S1. Statistics and analyses of sequencing data.**

**Supplementary Table S2. Relative abundance of fungi in bulk soil.**

**Supplementary Table S3. Relative abundance of fungi in rhizosphere soil.**

**Supplementary Table S4. Relative abundance increases multiples in rhizosphere fungal phyla compared with bulk soils.**

**Supplementary Table S5. OTU numbers and α-diversity of each sample.**

**Supplementary Table S6. Fungal genera that were increased or decreased in relative abundance in the rhizosphere compared with bulk soil in field soil.**

**Supplementary Table S7. Fungal genera that were increased or decreased in relative abundance in the rhizosphere compared with bulk soil in nutrient-rich soil.**

**Supplementary Table S8. Relative abundance of fungal genera that were affected by the presence of cotton root in nutrient-rich soil.**

**Supplementary Table S9. Relative abundance of genera that were affected by the presence of cotton root in field soil.**

**Supplementary Table S10. Analysis of fungal genera found during different plant developmental stages.**

**Supplementary Table S11. Beta-diversity between samples.**

**Supplementary Table S12. Analysis of potential pathogenic and phosphate-solubilizing fungal genera.**

## REFERENCES

1. Perez-Jaramillo JE, Mendes R, Raaijmakers JM. Impact of plant domestication on rhizosphere microbiome assembly and functions. Plant Mol Biol. 2016;90(6):635–44.

2. Wu Q-S, Zou Y-N, Huang Y-M. The arbuscular mycorrhizal fungus Diversispora spurca ameliorates effects of waterlogging on growth, root system architecture and antioxidant enzyme activities of citrus seedlings. Fungal Ecology. 2013;6(1):37–43.

3. Gannes Vd, Eudoxie G, Bekele I, Hickey WJ. Relations of microbiome characteristics to edaphic properties of tropical soils from Trinidad. Front Microbiol. 2015;6:1045.

4. Xu Z, Yu G, Zhang X, Ge J, He N, Wang Q, et al. The variations in soil microbial communities, enzyme activities and their relationships with soil organic matter decomposition along the northern slope of changbai mountain. Appl Soil Ecol. 2015;86:19–29.

5. Eva O, Barbara G, Wolfgang W, Andrea W, Christian S, Yvonne S, et al. Microbial decomposition of 13C-labeled phytosiderophores in the rhizosphere of wheat: Mineralization dynamics and key microbial groups involved. Soil Biol Biochem. 2016;98:196–207.

6. Itoh K. Study of the ecology of pesticide-degrading microorganisms in soil and an assessment of pesticide effects on the ecosystem. J Pestic Sci. 2014;39(3):174–6.

7. Kotoky R, Rajkumari J, Pandey P. The rhizosphere microbiome: Significance in rhizoremediation of polyaromatic hydrocarbon contaminated soil. Journal of Environmental Management. 2018;217:858–70.

8. Trivedi P, Delgado-Baquerizo M, Trivedi C, Hu H, Anderson IC, Jeffries TC, et al. Microbial regulation of the soil carbon cycle: evidence from gene-enzyme relationships. ISME J. 2016;10(11):2593–604.

9. Thion CE, Poirel JD, Cornulier T, De Vries FT, Bardgett RD, Prosser JI. Plant nitrogen-use strategy as a driver of rhizosphere archaeal and bacterial ammonia oxidiser abundance. FEMS Microbiol Ecol. 2016;92(7).

10. Cotta SR, Dias ACF, Seldin L, Andreote FD, Elsas JDv. The diversity and abundance of phytase genes (β-propeller phytases) in bacterial communities of the maize rhizosphere. Lett Appl Microbiol. 2016;62(3):264–8.

11. Kertesz MA, Mirleau P. The role of soil microbes in plant sulphur nutrition. J Exp Bot. 2004;55(404):1939.

12. Igiehon NO, Babalola OO. Rhizosphere Microbiome Modulators: Contributions of Nitrogen Fixing Bacteria towards Sustainable Agriculture. International Journal of Environmental Research & Public Health. 2018;15(4).

13. Ellouze W, Esmaeili Taheri A, Bainard LD, Yang C, Bazghaleh N, Navarro-Borrell A, et al. Soil Fungal Resources in Annual Cropping Systems and Their Potential for Management. BioMed Res Int. 2014;2014:15.

14. Wakelin SA, Warren RA, Harvey PR, Ryder MH. Phosphate solubilization by Penicillium spp. closely associated with wheat roots. Biol Fert Soils. 2004;40(1):36–43.

15. Mittal V, Singh O, Nayyar H, Kaur J, Tewari R. Stimulatory effect of phosphate-solubilizing fungal strains (*Aspergillus awamori* and *Penicillium citrinum*) on the yield of chickpea (*Cicer arietinum* L. cv. GPF2). Soil Biol Biochem. 2008;40(3):718–27.

16. Xiao C, Chi R, He H, Qiu G, Wang D, Zhang W. Isolation of phosphate-solubilizing fungi from phosphate mines and their effect on wheat seedling growth. Appl Biochem Biotechnol. 2009;159(2):330–42.

17. Kowalchuk GA, Veen JAHv. The significance of microbial diversity in agricultural soil for suppressiveness of plant diseases and nutrient retention. Physiol Behav. 2004;100(5):519–24.

18. Chapelle E, Mendes R, Bakker PA, Raaijmakers JM. Fungal invasion of the rhizosphere microbiome. ISME J. 2016;10(1):265–8.

19. Zhang Y, He J, Jia L-J, Yuan T-L, Zhang D, Guo Y, et al. Cellular tracking and gene profiling of fusarium graminearum during maize stalk rot disease development elucidates its strategies in confronting phosphorus limitation in the host apoplast. PLOS Pathog. 2016;12(3):e1005485.

20. Rebbeck J, Malone MA, Short D, Kasson MT, O’Neal ES, Davis DD. First report of verticillium wilt caused by *Verticillium nonalfalfaeon tree-of-heaven (Ailanthus altissima*) in Ohio. Plant Dis. 2013;97(7):999–1000.

21. Tetali S, Karpagavalli S, Pavani SL. Management of dry root rot of blackgram caused by *Macrophomina phaseolina* (Tassi) Goid. using bio agent. Plant Arch. 2015;15(2):647–50.

22. Garbeva P, Veen JAV, Elsas JDV. MICROBIAL DIVERSITY IN SOIL: Selection of Microbial Populations by Plant and Soil Type and Implications for Disease Suppressiveness. Annual Review of Phytopathology. 2004;42(42):243–70.

23. Chapelle E, Mendes R, Bakker PAH, Raaijmakers JM. Fungal invasion of the rhizosphere microbiome. Isme Journal. 2016;10(1):265–8.

24. Kowalchuk GA, Veen JAV. The significance of microbial diversity in agricultural soil for suppressiveness of plant diseases and nutrient retention. Physiology & Behavior. 2004;100(5):519–24.

25. Zhang Q, Gao X, Ren Y, Ding X, Qiu J, Li N, et al. Improvement of Verticillium Wilt Resistance by Applying Arbuscular Mycorrhizal Fungi to a Cotton Variety with High Symbiotic Efficiency under Field Conditions. International Journal of Molecular Sciences. 2018;19(1):241.

26. Tkacz A, Cheema J, Chandra G, Grant A, Poole PS. Stability and succession of the rhizosphere microbiota depends upon plant type and soil composition. ISME J. 2015;9(11):2349–59.

27. Kazeeroni EA, Al-Sadi AM. 454-pyrosequencing reveals variable fungal diversity across farming systems. Front Plant Sci. 2016;7:314.

28. Zarraonaindia I, Owens SM, Weisenhorn P, West K, Hampton-Marcell J, Lax S, et al. The soil microbiome influences grapevine-associated microbiota. mBio. 2015;6(2):e02527-14.

29. Bulgarelli D, Garrido-Oter R, Münch Philipp C, Weiman A, Dröge J, Pan Y, et al. Structure and function of the bacterial root microbiota in wild and domesticated barley. Cell host & microbe. 2015;17(3):392–403.

30. Shakya M, Gottel N, Castro H, Yang ZK, Gunter L, Labbe J, et al. A multifactor analysis of fungal and bacterial community structure in the root microbiome of mature Populus deltoides trees. PloS one. 2013;8(10):e76382.

31. Zhang W, Long X, Huo X, Chen Y, Lou K. 16S rRNA-Based PCR-DGGE Analysis of Actinomycete Communities in Fields with Continuous Cotton Cropping in Xinjiang, China. Microbial Ecol. 2013;66(2):385–93.

32. Vargas Gil S, Meriles J, Conforto C, Figoni G, Basanta M, Lovera E, et al. Field assessment of soil biological and chemical quality in response to crop management practices. World J Microbiol Biotech. 2009;25(3):439–48.

33. Peralta AL, Sun Y, Mcdaniel MD, Lennon JT. Crop rotational diversity increases disease suppressive capacity of soil microbiomes. Ecosphere. 2018;9(5):e02235.

34. Bai L, Cui J, Jie W, Cai B. Analysis of the community compositions of rhizosphere fungi in soybeans continuous cropping fields. Microbiol Res. 2015;180(Supplement C):49–56.

35. Fu Q, Liu C, Ding N, Lin Y, Guo B, Luo J, et al. Soil microbial communities and enzyme activities in a reclaimed coastal soil chronosequence under rice–barley cropping. J Soil Sediment. 2012;12(7):1134–44.

36. Bacharis C, Gouziotis A, Kalogeropoulou P, Koutita O, Tzavella-Klonari K, Karaoglanidis GS. Characterization of *Rhizoctonia* spp. isolates associated with damping-off disease in cotton and tobacco seedlings in Greece. Plant Dis. 2010;94(11):1314–22.

37. Sanogo S, Zhang J. Resistance sources, resistance screening techniques and disease management for Fusarium wilt in cotton. Euphytica. 2015;207(2):255–71.

38. Laidou IA, Koulakiotu EK, Thanassoulopoulos CC. First report of stem canker caused by Alternaria alternata on cotton. Plant Dis. 2007;84(1):103-.

39. Zhang W, Zhang H, Qi F, Jian G. Generation of transcriptome profiling and gene functional analysis in *Gossypium hirsutum* upon *Verticillium dahliae* infection. Biochem Bioph Res Co. 2016;473(4):879–85.

40. Knox O, Vadakattu G, Lardner R. Field evaluation of the effects of cotton variety and GM status on rhizosphere microbial diversity and function in Australian soils. Soil Res. 2014;52(2):203.

41. Lundberg DS, Lebeis SL, Paredes SH, Yourstone S, Gehring J, Malfatti S, et al. Defining the core Arabidopsis thaliana root microbiome. Nature. 2012;488(7409):86–90.

42. Qiao Q, Wang F, Zhang J, Chen Y, Zhang C, Liu G, et al. The variation in the rhizosphere microbiome of cotton with soil type, genotype and developmental stage. Sci Rep. 2017;7(1):3940.

43. Desjardins AE. Gibberella from A (venaceae) to Z (eae). Annu Rev Phytopathol. 2003;41:177–98.

44. Flentje N, Dodman R, Kerr A. The Mechanism of Host Penetration by Thanatephorus Cucumeris. Aus J Biol Sci. 1963;16(4):784–99.

45. Whalley WR, Riseley B, Leeds-Harrison PB, Bird NRA, Leech PK, Adderley WP. Structural differences between bulk and rhizosphere soil. Eur J Soil Sci. 2005;56(3):353–60.

46. Gould IJ, Quinton JN, Weigelt A, Deyn GBD, Bardgett RD. Plant diversity and root traits benefit physical properties key to soil function in grasslands. Ecol Lett. 2016;19(9):1140–39.

47. Odell RE, Dumlao MR, Samar D, Silk WK. Stage-dependent border cell and carbon flow from roots to rhizosphere. American journal of botany. 2008;95(4):441–6.

48. Bjørnlund L, Mørk S, Vestergård M, Rønn R. Trophic interactions between rhizosphere bacteria and bacterial feeders influenced by phosphate and aphids in barley. Biol Fert Soils 2006;43(1):1–11.

49. Stumpf L, Pauletto EA, Pinto LFS. Soil aggregation and root growth of perennial grasses in a constructed clay minesoil. Soil Till Res. 2016;161:71–8.

50. Zhu S, Vivanco JM, Manter DK. Nitrogen fertilizer rate affects root exudation, the rhizosphere microbiome and nitrogen-use-efficiency of maize. Appl Soil Ecol. 2016;107:324–33.

51. Watson BS, Bedair MF, Urbanczyk-Wochniak E, Huhman DV, Yang DS, Allen SN, et al. Integrated metabolomics and transcriptomics reveal enhanced specialized metabolism in *Medicago truncatula* root border cells. Plant Physiol. 2015;167(4):1699–716.

52. Haichar FeZ, Santaella C, Heulin T, Achouak W. Root exudates mediated interactions belowground. Soil Biol Biochem. 2014;77:69–80.

53. Huang XF, Chaparro JM, Reardon KF, Zhang R, Shen Q, Vivanco JM. Rhizosphere interactions: root exudates, microbes, and microbial communities1. Botany. 2014;92(4):267–75.

54. Plancot B, Santaella C, Jaber R, Kiefer-Meyer MC, Follet-Gueye M-L, Leprince J, et al. Deciphering the responses of root border-like cells of Arabidopsis and flax to pathogen-derived elicitors. Plant Physiol. 2013;163(4):1584.

55. Curlango-Rivera G, Huskey DA, Mostafa A, Kessler JO, Xiong Z, Hawes MC. Intraspecies variation in cotton border cell production: rhizosphere microbiome implications. Am J Bot. 2013;100(9):1706–12.

56. Kawasaki A, Donn S, Ryan PR, Mathesius U, Devilla R, Jones A, et al. Microbiome and exudates of the root and rhizosphere of brachypodium distachyon, a model for wheat. PloS one. 2016;11(10):e0164533.

57. Chen Z, Tian Y, Zhang Y, Song BR, Li H, Chen Z. Effects of root organic exudates on rhizosphere microbes and nutrient removal in the constructed wetlands. Ecol Eng. 2016;92:243–50.

58. Bulgarelli D, Schlaeppi K, Spaepen S, Themaat EVLv, Schulze-Lefert P. Structure and functions of the bacterial microbiota of plants. Annu Rev Plant Biol. 2013;64(1):807–38.

59. Edwardsa J, Johnsona C, Santos-Medellína C, Luriea E, Podishettyb NK, Bhatnagarc S, et al. Structure, variation, and assembly of the root-associated microbiomes of rice. Proc Nat Acad Sci. 2015;112(8):E911–E20.

60. Xu L, Ravnskov S, Larsen J, Nilsson RH, Nicolaisen M. Soil fungal community structure along a soil health gradient in pea fields examined using deep amplicon sequencing. Soil Biol Biochem. 2012;46:26–32.

61. Bakker MG, Chaparro JM, Manter DK, Vivanco JM. Impacts of bulk soil microbial community structure on rhizosphere microbiomes of Zea mays. Plant Soil. 2015;392(1-2):115–26.

62. Ling N, Kaiying D, Song Y, Wu Y, Zhao J, Raza W, et al. Variation of rhizosphere bacterial community in watermelon continuous mono-cropping soil by long-term application of a novel bioorganic fertilizer. Microbiol Res. 2014;169(7-8):570.

63. Gleń-Karolczyk K, Boligłowa E, Antonkiewicz J. Organic fertilization shapes the biodiversity of fungal communities associated with potato dry rot. Applied Soil Ecology. 2018.

64. Baudoin E, Benizri E, Guckert A. Impact of growth stage on the bacterial community structure along maize roots, as determined by metabolic and genetic fingerprinting. ApplSoil Ecol. 2002;19(2):135–45.

65. Okubo T, Tokida T, Ikeda S, Bao Z, Tago K, Hayatsu M, et al. Effects of elevated carbon dioxide, elevated temperature, and rice growth stage on the community structure of rice root-associated bacteria. Microbes Environ. 2014;29(2):184–90.

66. İnceoğlu Ö, Salles JF, Overbeek Lv, Elsas JDv. Effects of plant genotype and growth stage on the betaproteobacterial communities associated with different potato cultivars in two fields. Appl Environ Microbiol. 2010;76(11):3675–584.

67. Li X, Rui J, Mao Y, Yannarell A, Mackie R. Dynamics of the bacterial community structure in the rhizosphere of a maize cultivar. Soil Biol Biochem. 2014;68:392–401.

68. Breidenbach B, Pump J, Dumont MG. Microbial community structure in the rhizosphere of rice plants. Front Microbiol. 2016;6:1537.

69. Schlemper TR, Mfa L, Lucheta AR, Shimels M, Bouwmeester HJ, van Veen JA, et al. Rhizobacterial community structure differences among sorghum cultivars in different growth stages and soils. FEMS microbiology ecology. 2017;93(8):1–11.

70. Zhang T, Wang N-F, Liu H-Y, Zhang Y-Q, Yu L-Y. Soil pH is a key determinant of soil fungal community composition in the Ny-Alesund region, Svalbard (High Arctic). Front Microbiol. 2016;7:244.

71. Rahman M, Punja ZK. Factors Influencing Development of Root Rot on Ginseng Caused by Cylindrocarpon destructans. Phytopathology. 2005;95(12):1381–90.

72. Silva-Hughes AF, Wedge DE, Cantrell CL, Carvalho CR, Pan Z, Moraes RM, et al. Diversity and antifungal activity of the endophytic fungi associated with the native medicinal cactus Opuntia humifusa (Cactaceae) from the United States. Microbiol Res. 2015;175:67–77.

73. Silva GH, de Oliveira CM, Teles HL, Pauletti PM, Castro-Gamboa I, Silva DHS, et al. Sesquiterpenes from Xylaria sp., an endophytic fungus associated with Piper aduncum (Piperaceae). Phytochem Lett. 2010;3(3):164–7.

74. Zhang Y, Chen FS, Wu XQ, Luan FG, Zhang LP, Fang XM, et al. Isolation and characterization of two phosphate-solubilizing fungi from rhizosphere soil of moso bamboo and their functional capacities when exposed to different phosphorus sources and pH environments. PloS one. 2018;13(7):e0199625.

75. Reddy MS, Kumar S, Babita K. Biosolubilization of poorly soluble rock phosphates by Aspergillus tubingensis and Aspergillus niger. Bioresource Technology. 2002;84(2):187–9.

76. Morales A, Marysol A, Valenzuela E, Rubio R, Borie F. Effect of inoculation with *Penicillium albidum*, a phosphate-solubilizing fungus, on the growth of Trifolium pratense cropped in a volcanic soil. J Basic Microb. 2007;47(3):275–80.

77. Gong M, Du P, Liu X, Zhu C. Transformation of Inorganic P Fractions of Soil and Plant Growth Promotion by Phosphate-solubilizing Ability of Penicillium oxalicum I1. J Microbiol. 2014;52(12):1012–9.

